# ImmunoPET-informed sequence for focused ultrasound-targeted mCD47 blockade controls glioma

**DOI:** 10.1101/2020.07.27.223206

**Authors:** Natasha D. Sheybani, Soumen Paul, Katelyenn S. McCauley, Victoria R. Breza, Stuart S. Berr, G. Wilson Miller, Kiel D. Neumann, Richard J. Price

**Author notes:** Corresponding Author: Natasha D. Sheybani, Department of Biomedical Engineering, Box 800759, UVA Health System, University of Virginia, Charlottesville, VA 22908.

## Abstract

Phagocytic immunotherapies such as CD47 blockade have emerged as promising strategies for glioblastoma (GB) therapy, but the blood brain/tumor barriers (BBB/BTB) pose a persistent challenge for mCD47 delivery that can be overcome by focused ultrasound (FUS)-mediated BBB/BTB disruption. We here leverage immuno-PET imaging to determine how timing of [^89^Zr]-mCD47 injection relative to FUS impacts antibody penetrance into orthotopic murine gliomas. We then design and implement a rational paradigm for combining FUS and mCD47 for glioma therapy. We demonstrate that timing of antibody injection relative to FUS BBB/BTB disruption is a critical determinant of mCD47 access, with post-FUS injection conferring superlative antibody delivery to gliomas. We also show that mCD47 delivery across the BBB/BTB with repeat sessions of FUS can significantly constrain tumor outgrowth and extend survival in glioma-bearing mice. This study generates provocative insights for ongoing pre-clinical and clinical evaluations of FUS-mediated antibody delivery to brain tumors. Moreover, our results confirm that mCD47 delivery with FUS is a promising therapeutic strategy for GB therapy.

## Background

The natural disease course for glioblastoma (GB), the most common and aggressive primary cancer of the brain, is invariably grim. Even with the current standard treatment consisting of maximal safe resection, followed by irradiation and temozolomide chemotherapy, patients are faced with swift and devastating median survival outcomes on the order of 18 months [1,2]. Recent advances in cancer immunotherapy have reinvigorated hope for improved outcomes in GB. Targeting of phagocytosis checkpoints, such as the CD47–signal regulatory protein-α (SIRPα) axis, represents a promising new frontier for innate immune therapy approaches with demonstrated efficacy in many cancers including GB [3–5].

CD47 is ubiquitously overexpressed on the surface of most cancer cells [6]; its interaction with SIRPα on phagocytic cell types (e.g. macrophages) relays a “don’t eat me” signal that inhibits the activity of these cells as a mechanism of immune evasion [7,8]. Most healthy cells, including those in the CNS [9], do not express CD47 at appreciable levels [10]. As such, CD47 blockade via mCD47 shows some efficacy against GB [3,11]. However, while mCD47 has been shown to perform well in immunocompromised brain tumor-bearing mice, the outcomes have been far weaker in immunocompetent hosts [3]. Given what is known about how significantly the blood-brain and blood-tumor barriers (BBB/BTB) hinder antibody delivery [12,13], it is expected that lifting these barriers can improve mCD47 access – and consequently, outcomes of this therapy – in immunocompetent brain-tumor bearing animals.

The BBB is a highly exclusionary barrier that not only prevents the transport of large molecules such as antibodies in healthy brain tissue, but also protects the infiltrating rim of GB tumors from exposure to systemically administered chemo- and immunotherapies [14,15]. Meanwhile, the heterogeneously leaky vasculature of the BTB gives way to the “enhanced permeability and retention” (EPR) effect for passive drug transport; in this region, high interstitial fluid pressures can hinder convective transport and thereby penetrance of circulating therapies into tissues [16]. Focused ultrasound (FUS) in the presence of circulating microbubbles (MB) has gained remarkable traction as a non-invasive, non-ionizing technique for localized BBB/BTB disruption (BBB/BTB-D). Having been shown to circumvent these barriers safely and repeatably, FUS can confer meaningful delivery of drugs, genes and immunotherapies to a wide array of CNS pathologies, including GB [17]. For instance, studies have shown that FUS can improve outcomes in the context of antibody delivery to brain tumors, whether by systemic administration of antibody directly before FUS (e.g. trastuzumab [18], TDM1 [19]) or by administration following sonications (e.g. bevacizumab [20], trastuzumab [21]). While these studies lend to the promise of FUS for antibody delivery, they collectively highlight timing of antibody injection as a discrepant – and thus poorly understood – parameter among FUS BBB opening studies.

To date, no studies have delivered mCD47 across the BBB/BTB with FUS. In order to arrive at a rational protocol for combining FUS and mCD47 for GB therapy, we labeled a CD47-targeted antibody with zirconium-89 and leveraged Positron Emission Tomography (PET) to non-invasively profile the impact of FUS BBB/BTB-D on [^89^Zr]-mCD47 kinetics in the context of an orthotopic murine glioma model. We test the hypothesis that timing of injection relative to FUS impacts [^89^Zr]-mCD47 penetrance and retention in gliomas. Finally, we evaluate whether mCD47 delivery across the BBB/BTB-D with FUS offers protection against glioma in immunocompetent mice.

## Results

### FUS-mediated BBB/BTB disruption in murine GL261 gliomas

An orthotopic syngeneic glioma model was established via stereotaxic implantation of GL261 cells into the right striatum of C57Bl6 mice. Two weeks following intracranial tumor implantation, mice were randomized into three experimental groups, two of which underwent FUS-mediated BBB/BTB-D. MR-visible GL261 tumors were targeted with a 4-spot grid of sonications (1.1 MHz, 0.5% duty cycle, 0.4 MPa, 2 m duration) in the presence of systemically circulating albumin-shelled MB. At the conclusion of the two-minute sonication period, contrast-enhanced T1 weighted MR imaging was repeated to verify BBB/BTB-D (Figure 1A). Quantification of changes in mean greyscale intensity of MRI contrast enhancement revealed a significant increase in contrast enhancement from pre-FUS to post-FUS MR imaging, suggesting that the BBB/BTB was indeed effectively disrupted in and around GL261 tumors following FUS (Figure 1B).

**Figure 1.**
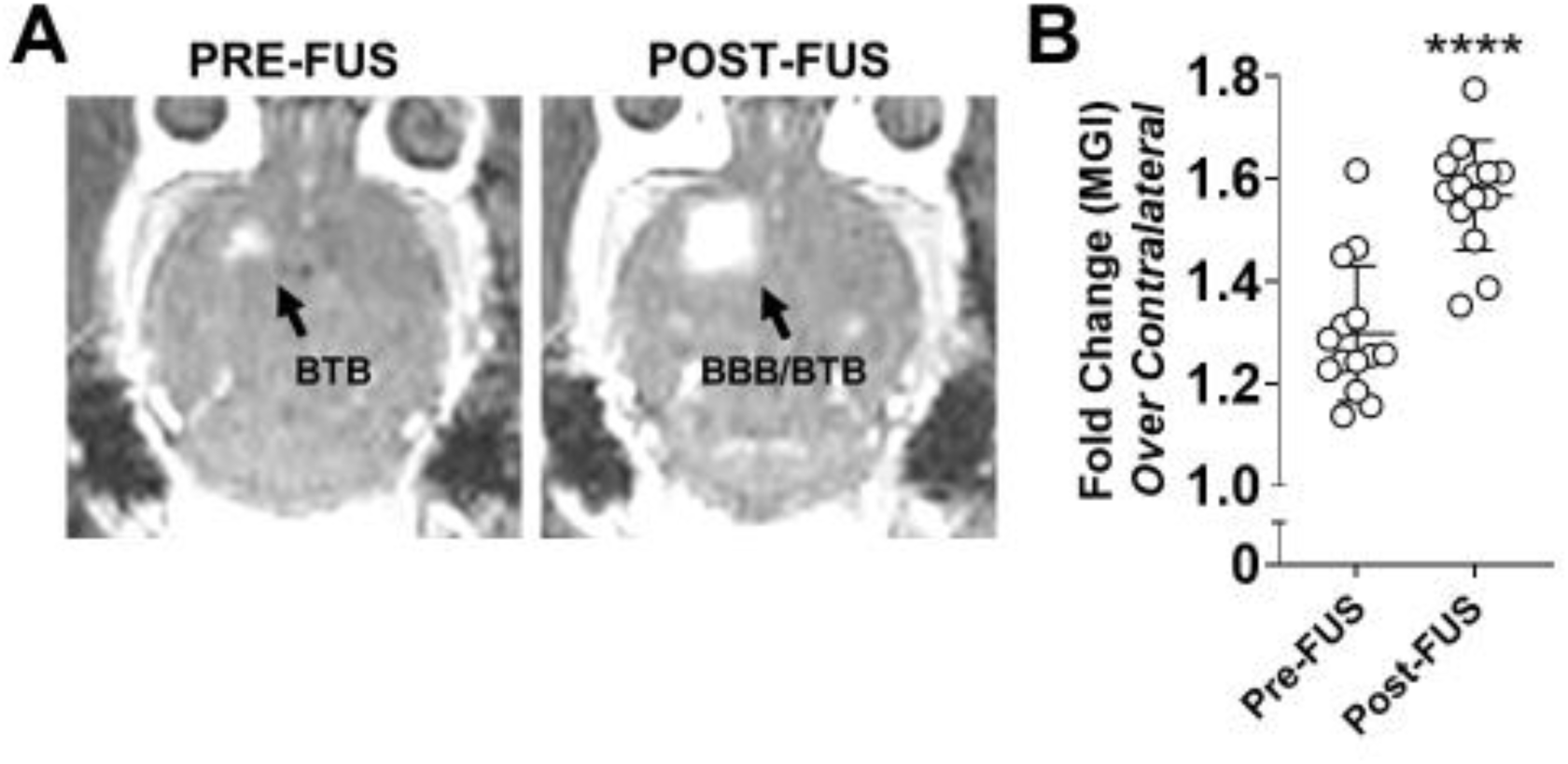
MR-image guided FUS-mediated blood brain/tumor barrier disruption in murine GL261 tumors. (A) Representative contrast-enhanced T1-weighted MR images of a GL261 tumor-bearing brain pre- and post-FUS. Enhancement in the right cerebral hemisphere on pre-FUS imaging indicates baseline barrier disruption induced by the presence of a brain tumor. Expansion of the enhanced region on post-FUS imaging reflects effective FUS-mediated BBB/BTB disruption. (B) Mean greyscale intensity (MGI) of contrast enhancement pre- and post-FUS BBB/BTB-D. Calculated as fold change over contralateral brain. Mean ± SD. ****p<0.0001. Significance assessed by unpaired t-test.

### Sequencing of [^89^Zr]-mCD47 delivery relative to MRI-guided FUS BBB/BTB-D is a critical determinant of antibody penetrance into murine gliomas

On day 14 post-implantation, GL261-bearing mice received either [^89^Zr]-mCD47 only [GL261 (EPR)], [^89^Zr]-mCD47 immediately followed by FUS BBB/BTB-D [GL261 + FUS_PRE_] or [^89^Zr]-mCD47 approximately 15 minutes following FUS BBB/BTB-D [GL261 + FUS_POST_]. In order to evaluate the hypothesis that mCD47 injection timing influences its accessibility to FUS-exposed tumors, we performed serial PET/CT imaging on [^89^Zr]-mCD47-recipient mice between 0 and 48 hours following (i.v.) injection and additionally assessed *ex vivo* biodistribution by automated gamma counter at a terminal time point (72 hours) (Figure 2). We performed an *in vitro* cell binding assay to confirm specificity of the [^89^Zr]-mCD47 formulation for CD47 enriched on the surface of GL261 cells. Our results confirmed an average (± SD) of 11.22 ± 0.86% cell associated activity.

**Figure 2.**
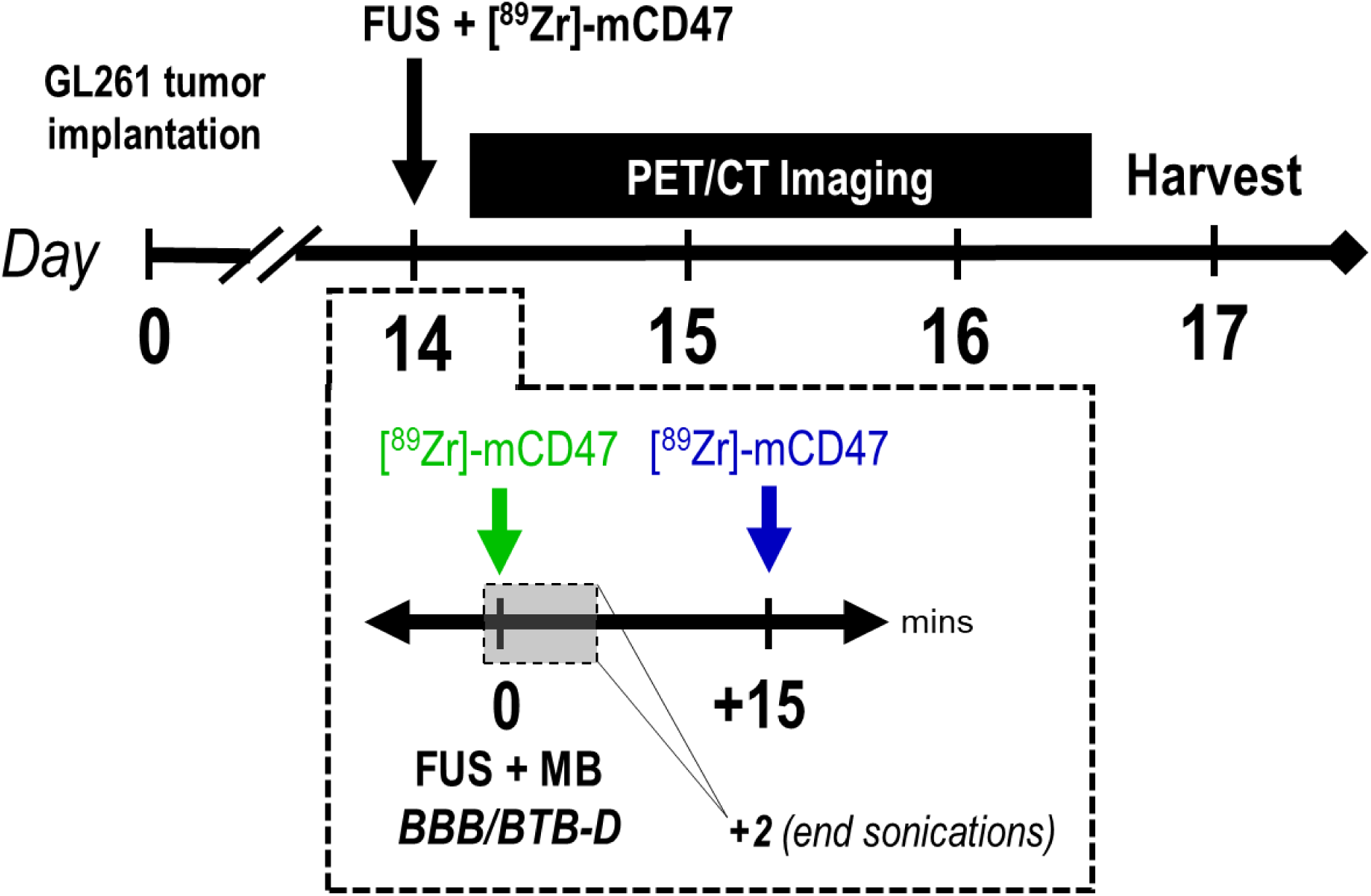
Overview of integrated imaging workflow for interrogating impact of FUS BBB/BTB-D on [89Zr]-mCD47 delivery. Two weeks following intracranial implantation of GL261 tumors, mice were randomized into three groups receiving either [89Zr]-mCD47 only [GL261 (EPR)], [89Zr]-mCD47 immediately followed by FUS BBB/BTB-D [GL261 + FUS_PRE_; indicated in green] or [89Zr]-mCD47 approximately 15 minutes following FUS BBB/BTB-D [GL261 + FUS_POST_; indicated in blue]. Mice subsequently underwent serial static PET/CT imaging (on Days 14, 15, 16) and terminal biodistribution assessment by automated gamma counter (on Day 17).

Though MR imaging confirmed the presence of a tumor in all mice, PET/CT imaging did not reveal clear preferential [^89^Zr]-mCD47 uptake at the site of GL261 tumors via EPR or BBB/BTB-D with the FUS_PRE_ protocol. Remarkably, however, FUS_POST_ conferred superlative [^89^Zr]-mCD47 uptake at the site of BBB/BTB-D relative to other groups (Figure 3A-B). This was evident both qualitatively in decay-corrected PET/CT images (Figure 3A-B; white arrows) as well as quantitatively from whole brain standardized uptake values (SUVs) extracted from these images (Figure 3C). Relative to all other experimental groups, [^89^Zr]-mCD47 uptake was significantly increased with implementation of the FUS_POST_ protocol. This trend was consistently observed between 0 and 48 hours post-FUS. [^89^Zr]-mCD47 uptake increased by ∼1.3-1.8-fold over naïve owing to the presence of a GL261 tumor (EPR effect only). Notably, the FUS_PRE_ protocol did not result in greater antibody penetrance over that established in the setting of control GL261 tumors. On day 14, uptake was significantly increased in the GL261 alone (p=0.0321) and GL261+FUS_PRE_ (p=0.0273) groups relative to naïve group. Remarkably, mice that underwent the FUS_POST_ protocol saw elevated [^89^Zr]-mCD47 uptake; 2.7- and 1.5-fold above their tumor-bearing and naïve counterparts, respectively. Interestingly, this stratification between groups grew over time, with the GL261+FUS_POST_ group boasting between 4.3- to 6.7-fold greater signal relative to other groups (Figure 3C). An additional statistical comparison of SUVs across imaging time points revealed that the temporal increase in uptake from Day 14 to 16 with FUS_POST_ was also significant (p<0.05 for all time point comparisons). Tumor-drug exposure was nearly 6- and 4-fold greater in GL261 tumors treated with FUS_POST_ relative to other GL261-bearing and naïve brains, respectively (Figure 3D).

**Figure 3.**
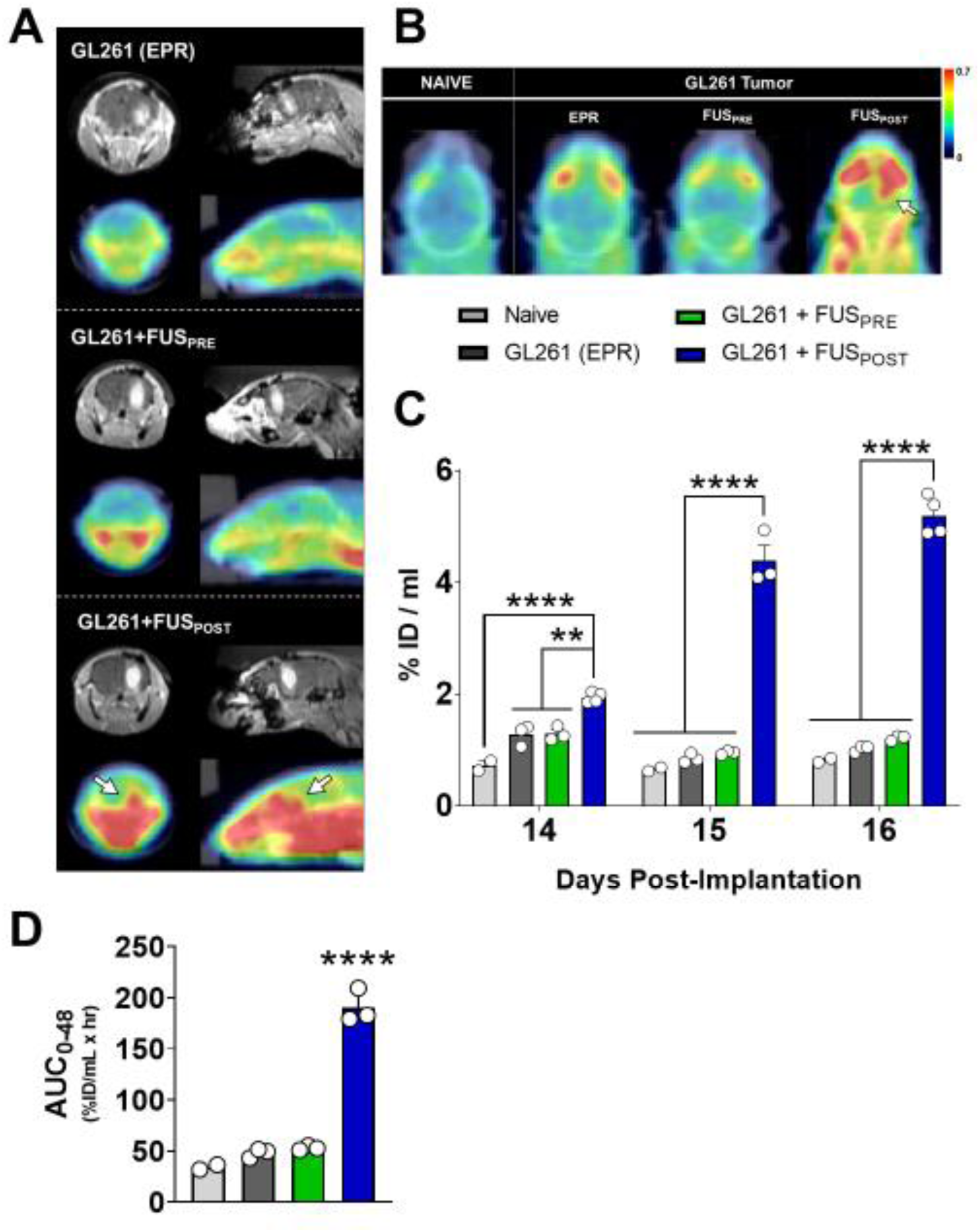
Sequencing of [89Zr]-mCD47 injection relative to FUS BBB/BTB-D markedly impacts antibody accumulation in murine GL261 gliomas. (A) Representative coronal (left) and sagittal (right) contrast-enhanced T1-weighted MR images (top row) and decay-corrected PET/CT images (bottom row) for each experimental group on Day 14. MR images depict a clear, enhancing tumor in the right hemisphere of GL261-bearing mice. (B) Representative axial decay-corrected PET/CT images for each experimental group on Day 14. White arrows denote region of visibly elevated radioactivity at the tumor site targeted with FUS. (C) Whole brain standardized [89Zr]-mCD47 uptake values (SUVs) extracted from serial static PET/CT images obtained between days 14 and 16 post-implantation (0 to 2 days post-[89Zr]-mCD47 injection). %ID/mL = % injected dose per mL. **p<0.01, ****p<0.0001 vs. group(s) indicated. Significance assessed by RM mixed effects model implementing restricted maximum likelihood method, followed by Tukey multiple comparison correction. (D) Tumor-drug exposure for [89Zr]-mCD47 in naïve brain or GL261 tumors, based on integration of SUVs from decay-corrected PET/CT images collected between 0 and 48 hours after BBB/BTB-D and/or [89Zr]-mCD47 injection. Significance assessed by one-way ANOVA followed by Tukey multiple comparison correction. ****p<0.0001 vs. all other groups.

Three days following FUS BBB/BTB-D and/or [^89^Zr]-mCD47 injection, we harvested brains (Figure 4A) and peripheral organs in order to assess [^89^Zr]-mCD47 biodistribution by automated gamma counter measurement. Consistent with our findings from PET/CT imaging, we observed that the FUS_POST_ protocol conferred significantly greater [^89^Zr]-mCD47 uptake in GL261 tumors relative to the baseline uptake within control tumors or naïve brains (Figure 4B). In the presence of a GL261 tumor, the right hemisphere of the brain also saw preferential uptake as expected (Figure 4C). The FUS_POST_ protocol most markedly enhanced this uptake, by 2.2- and 1.7-fold relative to control GL261 tumors and FUS_PRE_ protocol, respectively (Figure 4C). Accordingly, the percentage of whole brain radioactivity comprised by the right hemisphere was significantly higher in the GL261+FUS_POST_ group as well, once again suggesting preferential [^89^Zr]-mCD47 uptake in the tumor-bearing hemisphere. The mere presence of a GL261 tumor in the right hemisphere increased uptake by 35% relative to naïve brain. Relative to baseline uptake in GL261 tumors, the FUS_POST_ protocol then promoted a ∼21% increase in hemispheric [^89^Zr]-mCD47 uptake (Figure 4D). Overall, these findings, consistent with those unveiled by PET/CT imaging, confirmed the importance of antibody sequencing relative to FUS important.

**Figure 4.**
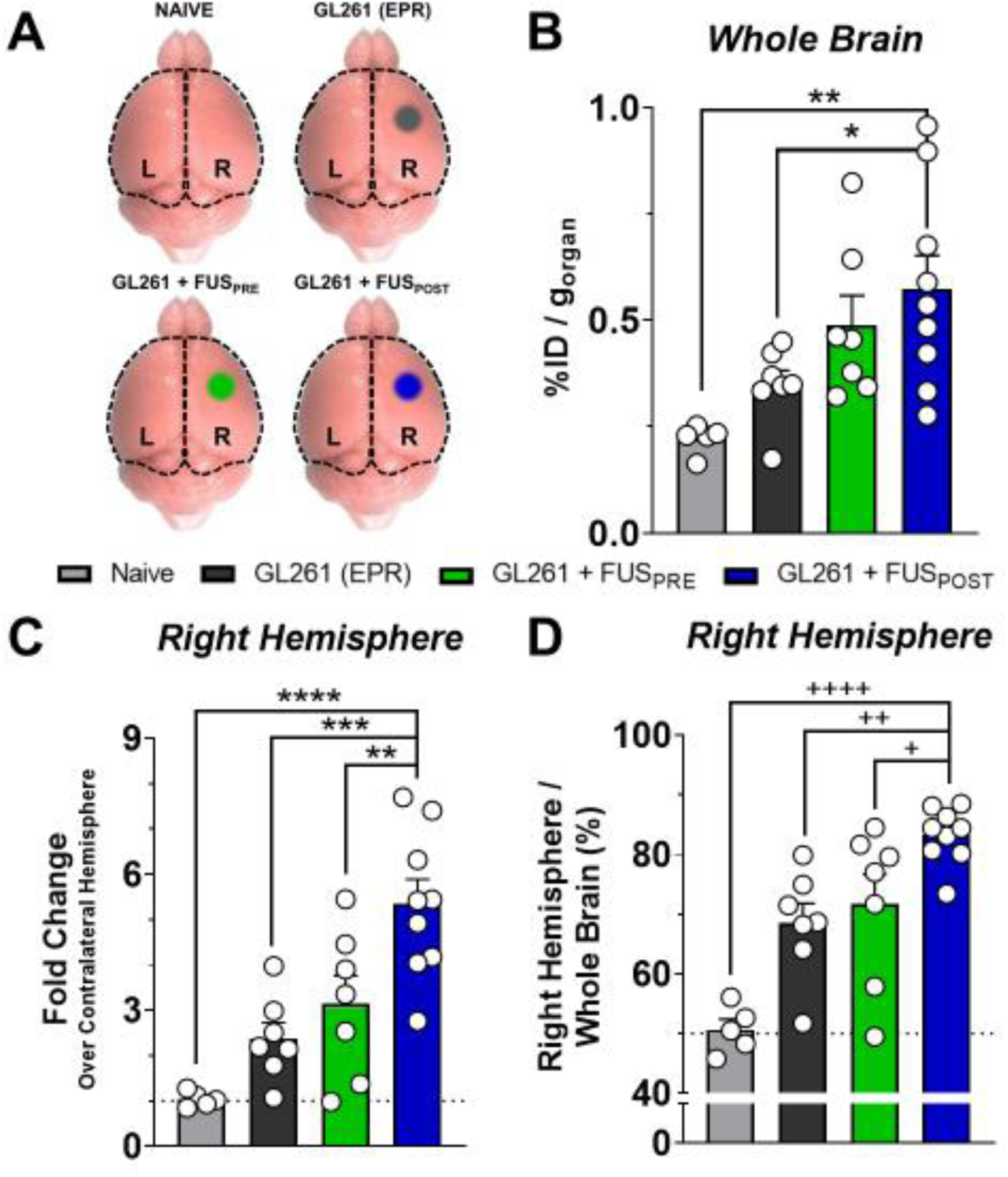
Ex vivo quantification of [89Zr]-mCD47 accumulation in naïve, tumor-bearing and FUS-treated animal brains by automated gamma counter measurement. (A) Schematic illustrating regions of brain harvested for automated gamma counter measurements from each experimental group. Right (R) and left (L) hemispheres of the cerebral cortex were separated at the midline and analyzed independently. Whole brain signal was quantified via summation of hemisphere signals (black dotted outline). (B) Whole brain accumulation of [89Zr]-mCD47 three days following i.v. injection and/or FUS BBB/BTB-D in naïve or tumor-bearing mice. *p=0.0430, **p=0.0031 vs. indicated group. (C-D) Right brain hemisphere accumulation of [89Zr]-mCD47 as fold change (%ID/g) over contralateral hemisphere (C) and as percentage of whole brain (D). **p=0.0086, ***p=0.0005, ****p<0.0001, ^+^p=0.0389, ^++^p=0.0065, ^++++^p<0.0001 vs. indicated group. Significance in all graphs assessed by One-way ANOVA followed by Sidak’s multiple comparison test.

### FUS BBB/BTB-D does not influence distribution of [89Zr]-mCD47 in the circulation or peripheral tissues of GL261-bearing mice

In tandem with evaluation of [^89^Zr]-mCD47 uptake in the brain by automated gamma counter, we collected peripheral blood and excised liver, spleen and kidneys in order to evaluate peripheral [^89^Zr]-mCD47 biodistribution across experimental groups. FUS BBB/BTB-D did not confer significant changes in [^89^Zr]-mCD47 signal within the circulation relative to control GL261 tumors (Figure 5A). Despite the marked changes noted in intracranial [^89^Zr]-mCD47 uptake following FUS_POST_, we noted that BBB/BTB-D did not influence biodistribution within the kidneys, liver or spleen of GL261-bearing mice (Figure 5B-E). Taken together, these observations suggest the localized influence of FUS BBB/BTB-D on antibody penetrance without consequence to the periphery.

**Figure 5.**
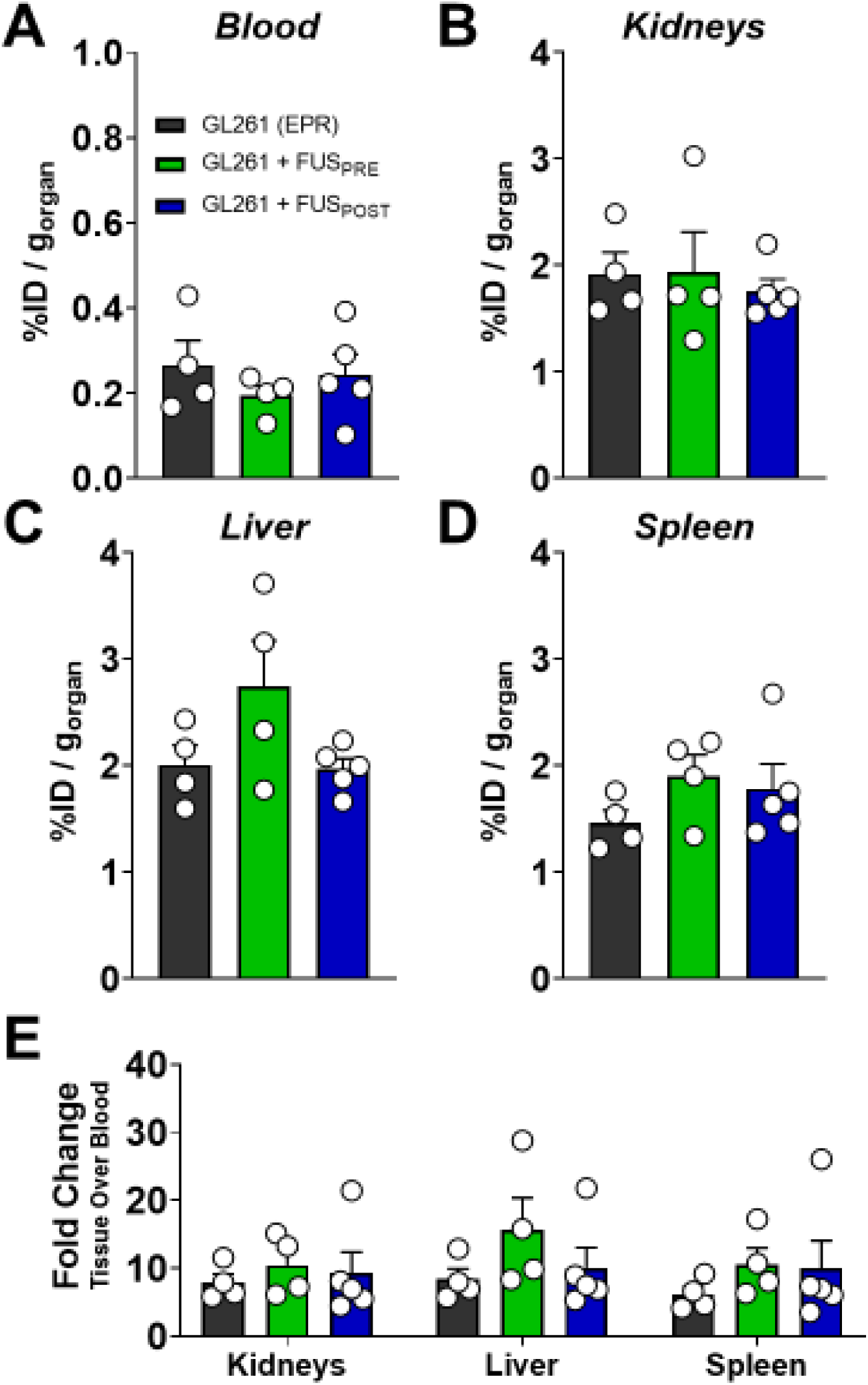
FUS BBB/BTB-D does not influence distribution of [89Zr]-mCD47 in the circulation or peripheral tissues of GL261-bearing mice. (A-D) [89Zr]-mCD47 distribution in circulation (A), kidneys (B), liver (C), and spleen (D) of GL261-bearing mice three days following systemic injection with or without FUS BBB/BTB-D. (E) Tissue to blood ratio calculated by normalizing [89Zr]-mCD47 distribution (%ID/g) in peripheral tissues to blood signal presented in (A). Significance in all graphs assessed by One-way ANOVA followed by Sidak’s multiple comparison test. No significant differences detected.

### Delivery of mCD47 via repeat FUS BBB/BTB-D significantly constrains tumor outgrowth and extends overall survival in glioma-bearing mice

Following on the insights we gleaned from immuno-PET imaging regarding mCD47 sequencing relative to BBB/BTB-D, we next designed and tested a rational therapeutic paradigm combining mCD47 with the FUS_POST_ protocol (Figure 6A). Beginning on Day 14 post-implantation, GL261-bearing mice underwent three sessions of repeat BBB/BTB-D every three days (Figure 6B-C). Control (FUS^-^) mice received mCD47 monotherapy (8 mg/kg, i.v.) while those undergoing BBB/BTB-D (FUS^+^) received mCD47 approximately 15 minutes following conclusion of sonications. Serial contrast-enhanced MR imaging revealed that FUS-mediated delivery of mCD47 across the BBB/BTB-D significantly constrained GL261 tumor outgrowth (Figure 6D). By day 20 post-implantation, we observed a 2.4-fold reduction in FUS^+^ tumor volume relative to FUS^-^ counterparts (Figure 6E). The protective effect of FUS-mediated mCD47 delivery was recapitulated in overall survival of GL261-bearing mice. The addition of BBB/BTB-D extended overall survival of mCD47-recipient mice by nearly 17% (hazard ratio: 0.2139) (Figure 6F). Importantly, whereas previous studies required a total mCD47 dose of >400 mg/kg to extend survival of GL261 tumor-bearing immunocompetent mice [3], we achieved this result using only 24 mg/kg due to immuno-PET-informed implementation of FUS BBB/BTB-D sequencing. Of note, in a separate cohort of mice, delivery of control IgG to GL261-bearing mice resulted in overall survival comparable to that observed in the FUS^-^ group (Supp. Figure 1).

**Figure 6.**
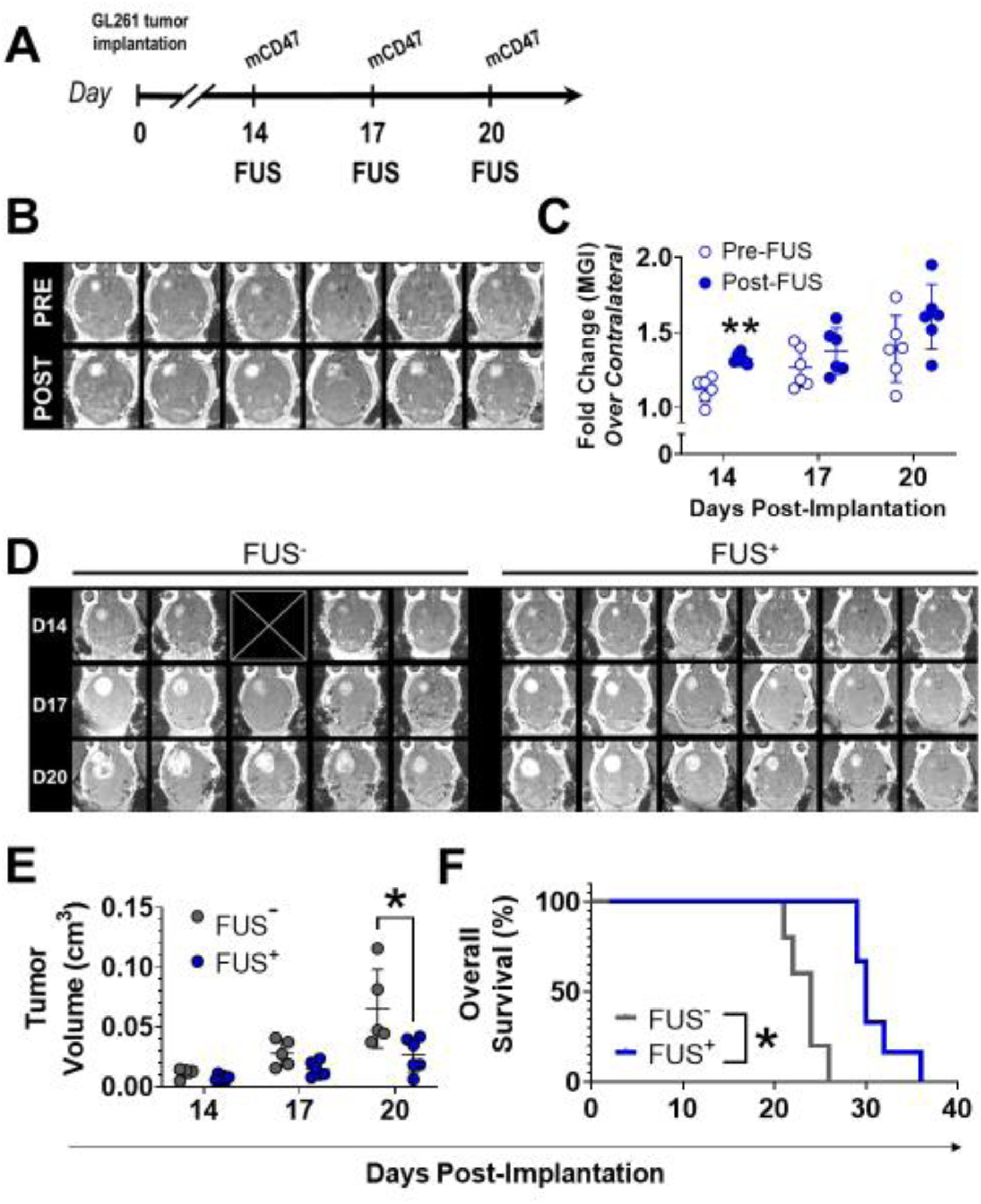
Delivery of mCD47 across the BBB/BTB with FUS significantly improves survival and restricts tumor outgrowth in GL261-bearing mice. (A) Overview of experimental design for evaluating therapeutic efficacy of mCD47 delivery to orthotopically implanted GL261 tumors in the context of repeat BBB/BTB-D with FUS. Mice received i.v. mCD47 (8mg/kg) either alone (FUS^-^) or following BBB/BTB-D at 0.4 MPa (FUS^+^). (B) Axial contrast-enhanced T1-weighted MR images of murine GL261-tumors pre- and post-FUS. (C) Mean greyscale intensity (MGI) of contrast enhancement pre- and post-FUS over three separate BBB/BTB-D sessions conducted every three days. Calculated as fold change over contralateral brain. Mean ± SD. **p=0.0023. Significance assessed by RM 2-way ANOVA followed by Sidak’s multiple comparison test. (D) Contrast-enhanced T1-weighted MR images of GL261 tumors on days 14, 17 and 20 post-implantation. ⊠ = image excluded due to poor quality. (E) GL261 tumor outgrowth quantified based on serial MR imaging. Mean ± SD. *p=0.0010. Significance assessed by RM mixed-effects model implementing restricted maximum likelihood method, followed by Sidak’s multiple comparison test. (F) Kaplan-Meier curve depicting overall survival of GL261-bearing mice. *n*=5-6 mice per group. *p=0.0008. Significance assessed by log-rank (Mantel-Cox) test.

## Discussion

In this study, we leveraged focused ultrasound BBB/BTB opening to improve cerebral access of a monoclonal antibody targeting the CD47-SIRPα axis in murine gliomas. Using an integrated MR-guided FUS and immuno-PET imaging workflow, we demonstrate that FUS BBB/BTB-D can play a profound role in improving mCD47 access within murine GL261 tumors, but only when mCD47 is systemically injected after FUS. With this finding, we demonstrate that injection timing relative to FUS - a parameter that has not been systematically examined within the literature to date - may be a critical determinant of antibody penetration into brain tumors. Using these insights, we designed a rational therapeutic paradigm combining repeated FUS BBB/BTB-D with post-FUS delivery of mCD47. In turn, this approach allowed us to control GL261 tumor growth and extend animal survival using an ∼20-fold lower total dosage of mCD47 compared to a previous study [3]. These findings constitute the first demonstration of a promising combinatorial therapy approach using mCD47 and FUS BBB/BTB-D. Taken together, our findings offer important and provocative insights for future pre-clinical and clinical investigations of FUS-mediated immunotherapy for brain tumors.

One of the goals of our study was to use non-invasive quantitative imaging for spatiotemporal mapping of monoclonal antibody delivery into GL261 tumors following FUS – as a prelude to combinatorial therapy paradigms utilizing antibody-based immunotherapies (e.g. checkpoint inhibitors). In order to limit confounding factors introduced by interactions of the antibody with systemic targets, we elected to use a cancer cell-specific therapeutic. Barring erythrocytes, CD47 is not expressed by normal cells [10]. Moreover, both murine GL261 tumors and human GBs have been demonstrated to richly overexpress CD47 relative to normal brain tissue [9,11]. Together, these considerations rendered mCD47 an excellent candidate for interrogating the influence of FUS on a tumor-targeted antibody.

Serial PET/CT imaging revealed that, despite the heterogeneously leaky vasculature evidenced by contrast enhancement of GL261 tumors on MRI, [^89^Zr]-mCD47 penetration into GL261 tumors was modest relative to naïve brain tissue. The less than 2-fold change observed in [^89^Zr]-mCD47 uptake following i.v. injection on Day 14 post-implantation was largely diminished by subsequent days of PET/CT imaging. This observation generates an important insight for myriad studies that are investigating the impact of mCD47 delivery in GL261 tumors, as it suggests that the enhanced permeation and retention (EPR) effect in these tumors is not very strong at baseline. Indeed, a recent study confirms that mCD47 (MIAP410) has limited anti-tumor effect when administered systemically in mice bearing orthotopically implanted syngeneic GL261 tumors [11]. In yet another study, GL261 tumors showed a seemingly dose-dependent survival response to mCD47; yet, complete eradication was not achieved even at the highest dose evaluated [3]. Despite the promise that these findings lend to mCD47 immunotherapy in gliomas, opportunities for improvement are limited as the direct mechanism of antibody transport through the BBB is yet unknown. Our results suggest that a persistent locoregional delivery challenge imposed by the BBB/BTB may be contributing to the marginal anti-glioma benefits observed in these studies.

We observed that contrast-enhanced MR imaging was poorly predictive of mCD47 penetrance into GL261 tumors. Indeed, in the GL261 (EPR) and GL261+FUS_PRE_ groups, where volumetric contrast enhancement was appreciable both before and even more so after FUS BBB/BTB-D, PET/CT imaging did not reveal similar spatial distribution of [^89^Zr]-mCD47. Quantitation from these images further revealed that the low [^89^Zr]-mCD47 uptake observed in these groups was conserved temporally. Given that a majority of clinical transcranial FUS studies are performed under the guidance of MRI and utilize contrast-enhanced imaging [22–24], the ability to use MR imaging as a surrogate for estimating drug delivery holds great appeal. Studies have related metrics from static or dynamic contrast-enhanced MR imaging to drug delivery in the context of small molecule chemotherapies on the order of 300-500 Da [25–27]. Good positive correlation has even been shown between trastuzumab delivery and MR-intensity following FUS BBB-D in normal brain tissue [28]. However, our findings suggest gadolinium leakage to be an inaccurate proxy in the case of mCD47 delivery in GL261 tumors. Multihance - the gadolinium-based contrast agent this study - has a molecular weight (MW) of 1.058 kDa, while most immunoglobulin G (IgG) antibodies have a MW on the order of 150 kDa. This stark difference in MW vastly underpins the false predictive value of MRI contrast enhancement that we observed in the context of mCD47 delivery and suggests that use of MRI-based changes should be coupled with more robust quantitative measures of effective delivery in the case of discrepantly sized drugs such as antibodies.

A wealth of studies in the literature have demonstrated that FUS BBB/BTB-D mediates antibody delivery to the brain in normal and diseased settings (e.g. brain tumors, Alzheimer’s disease) [20,28–34]. Among brain tumor studies in the literature, there exists a discrepancy in timing of antibody injection relative to FUS, with both protocols involving injection prior to [34] and following FUS [20] having shown therapeutic efficacy. While these studies have notably been executed in distinct models – brain metastasis of breast cancer and glioblastoma, respectively – they draw attention to a lack of consensus on optimal timing of therapeutic injection with respect to FUS BBB/BTB-D. Indeed, this is evident in the execution of clinical FUS BBB/BTB-D protocols in GB, where there is a similar disparity in choice of chemotherapy injection timing relative to FUS; some protocols perform systemic injection prior to sonication [24] and others do so after [35].

The dogmatic approach within the domain of FUS BBB/BTB-D has been to systemically inject therapeutic agents immediately prior to sonications [25,36–38]. We hypothesized that the FUS_PRE_ protocol would significantly increase [^89^Zr]-mCD47 uptake in GL261 tumors. Remarkably, FUS_PRE_ did not offer any significant benefit in antibody access over the control GL261 setting despite resulting in MR-visible BBB/BTB-D consistent with that observed in the FUS_POST_ protocol. Automated gamma counter measurements corroborated the results of PET/CT imaging. Owing to the greater sensitivity of this technique, we could appreciate a trend in [^89^Zr]-mCD47 intracerebral uptake from GL261 (EPR) to GL261+FUS_PRE_. However, this trend was not statistically significant. Other studies have evidenced that antibody delivery across the BBB/BTB with FUS_PRE_ is not always met with therapeutic response, an outcome for which our observations may offer an explanation [39]. Multiple incongruities between our study and others may explain the differences in performance of the FUS_PRE_ protocol - including MB formulation, anesthesia, transducer frequency, peak negative pressure (PNP), tumor model and targeted antibody. Of note, our BBB/BTB-D parameters (0.4 MPa PNP at 1.1 MHz) corresponded with a much lower mechanical index (MI = 0.38) relative to those of comparable tumor-targeted antibody delivery studies (MI = 0.6+) [20,29].

Our most striking observation was that a shift to post-FUS injection enabled the same FUS exposure conditions to reproducibly confer a marked increase in [^89^Zr]-mCD47 uptake. Serial PET/CT imaging revealed that this increased uptake was sustained well beyond the initial time point of intervention. Automated gamma counter measurements confirmed elevation of preferential uptake within the right hemisphere following the FUS_POST_ protocol. In the periphery, there were no appreciable changes in circulating [^89^Zr]-mCD47, nor were there changes in uptake across liver, kidneys and spleens of tumor-bearing mice. Taken together, these observations suggest that the effect of FUS_POST_ was localized with minimal off-target impact; elevated uptake was observed strictly in the CNS and FUS_POST_ did not pose a risk of elevated toxicity in the periphery.

Since immuno-PET imaging revealed FUS_POST_ to be the most effective regime for increasing antibody access in GL261 tumors, we translated this protocol into a treatment paradigm utilizing repeated BBB/BTB-D and a therapeutically relevant dose of mCD47. Barrier disruption was performed every three days in order to permit sufficient recovery time between sessions, and total sessions were limited to three owing to technical constraints on repeated tail vein cannulation. All mice received i.v. mCD47 at a dose of 8 mg/kg, with (FUS^+^) or without (FUS^-^) BBB/BTB-D. Combinatorial therapy markedly restricted tumor outgrowth by the final FUS session, with FUS+ GL261 tumors possessing a significantly lower total enhancing volume as compared with their monotherapy counterparts. Overall survival was consequently extended in FUS+ mice as well. Neither metric was reflective of complete tumor eradication, which is consistent with findings of other studies delivering this mCD47 clone in the GL261 model [3,11]. mCD47 has been previously shown to enhance survival of GL261-bearing mice in a dose-dependent manner with 16 mg/kg and 32 mg/kg systemic injections administered daily for a total of approximately 27 injections [3]. For the lower dose, this amounted to approximately 432 mg/kg of systemically administered antibody in total. Our results demonstrated that delivery of mCD47 across the BBB/BTB with FUS can effect the same degree of survival benefit, but with an ∼18-fold decrease in administered drug. This finding reveals the exceptional capacity of FUS to reduce the dose required to achieve therapeutic efficacy and mitigate risks of severe off-target toxicity associated, not only with mCD47 [40], but also with other immunotherapies as well [41].

In normal brain tissues, controlled FUS-mediated BBB opening has been estimated to last on the order of 4-6 hours [42]. Anecdotally, MR Imaging of human GBs following FUS BBB/BTB-D has shown recovery to baseline within 24 hours [24]. Here, serial PET/CT imaging revealed that intracerebral mCD47 uptake continued to increase significantly over the course of 48 hours subsequent to FUS_POST_-mediated delivery (Figure 3C). Given that this time frame surpasses the canonical time frame for BBB/BTB opening, we hypothesized that FUS BBB/BTB-D-mediated CD47 blockade could be sustaining BBB/BTB-D through a biological mechanism(s). Indeed, in addition to its demonstrated role in elaborating anti-tumor immune responses, blockade of the CD47-SIRPα axis has been shown to upregulate VEGF-A and VEGF-R2 expression, thereby promoting tumor permeability [43,44]. To test our hypothesis, we evaluated baseline permeability of GL261 tumors in our pre-treatment planning contrast MRIs from the mCD47 survival study. Here, we observed significant elevation of contrast enhancement in FUS^+^ tumors at both Day 17 and 20 (Supp. Figure 2), consistent with the hypothesis that high-levels of mCD47 sustain vessel permeabilization between FUS BBB/BTB-D sessions, leading to even further enhancement of total drug delivery through the entire experiment.

FUS-mediated barrier disruption has been widely regarded as a “drug neutral” method, owing to the diverse array of therapeutic agents it is capable of delivering across the BBB and/or BTB [45]. Our findings temper this argument with important considerations for agent-specific optimization of FUS-mediated drug delivery protocols. The urgency for systematic pre-clinical investigation of agent- or model-specific pharmacokinetics is further heightened by the pace of clinical activity; indeed, the first clinical trial delivering an immunotherapeutic antibody into brain tumors with FUS BBB/BTB-D is already underway (NCT04021420).

FUS BBB/BTB-D is a promising strategy for mCD47 delivery in GB. With the expanding role of nuclear medicine in FUS therapy [46], our findings lend credence to the utility of non-invasive techniques such as immuno-PET imaging for the design and monitoring of FUS immunotherapy paradigms. PET imaging has already been demonstrated to serve as a powerful tool for gaining robust insights into the mechanisms underlying FUS BBB disruption [22,47]. The workflow established within this study reiterates the tremendous and yet untapped potential of molecular imaging as an adjunct to FUS-mediated therapeutic paradigms [46].

Given the burgeoning role of BBB/BTB-D in clinical brain tumor care, these findings also generate compelling insights and important considerations for the delivery of immunotherapeutic antibodies with FUS, all the while expounding a novel paradigm for mCD47 therapy. Clinical development of CD47-targeting therapies is well underway [48], but targeting of CD47 within gliomas has been persistently challenged by presence of the BBB and BTB. Our readily translatable study generates timely evidence for the potential of MR image-guided FUS BBB/BTB-D to surmount this challenge and promote efficacy of mCD47 in the setting of GB.

## Methods

### Cell line and culture

Luciferase-transduced GL261 cells (GL261-luc2) obtained from the Woodworth Lab (University of Maryland) were cultured in high glucose 1x Dulbecco modified Eagle medium (DMEM, Gibco) supplemented with 1 mM sodium pyruvate (Gibco), non-essential amino acids (Gibco), 10% fetal bovine serum (Gibco), and 100 ug/mL G418 (GoldBio). Cells were maintained at 37°C and 5% CO_2_. Thawed cells were cultured for up to three passages and maintained in logarithmic growth phase for all experiments. Cells tested negative for mycoplasma.

### Intracranial tumor cell inoculation

GL261 cells (1 x10^5^ cells per 2µL) were resuspended in sterile PBS for intracranial tumor implantation. Cells were implanted into the right striatum of 6-10 week old female C57BL/6 mice (The Jackson Laboratory). Following anesthetization with an intraperitoneal injection of ketamine (50 mg/kg; Zoetis) and dexdomitor (0.25 mg/kg; Pfizer) in sterilized 0.9% saline, the heads of mice were depilated, aseptically prepared, and placed into a stereotactic head frame. Cells were implanted at a mechanically controlled rate of 0.5 μL/min using a Hamilton syringe and micropump (UltraMicroPump, World Precision Instruments). Injection coordinates were ∼2.0 mm lateral from the sagittal suture, 0.5mm anterior of bregma and 3mm below the dura. Mice were housed on a 12/12 h light/dark cycle and supplied food *ad libitum*. Animal experiments were approved by the Animal Care and Use Committee at the University of Virginia and conformed to the National Institutes of Health guidelines for the use of animals in research.

### MRI-guided focused ultrasound for single or repeat BBB/BTB disruption

Either 8 or 14 days following brain tumor inoculation, mice were anesthetized with isoflurane delivered to effect in concentrations of 2% in medical air using a vaporizer. Mice were maintained under anesthesia via nosecone throughout the FUS procedure. Tails were cannulated with a tail vein catheter to allow for intravenous (i.v.) injections of MRI contrast agent and microbubbles. Mouse heads were shaved and depilated in preparation for MRI-guided focused ultrasound treatment. Each mouse was positioned supine on a custom surface transmit-receive RF coil (to maximize imaging SNR) with the head coupled to a 1.14 MHz spherical single-element transducer via degassed water bath housed within an MR-compatible FUS system (RK-100; FUS Instruments). Image-guidance was enabled by co-registration of FUS system coordinates to those of either a 1.5T MRI (Avanto, Siemens) or 3T MRI (Prisma, Siemens) within which the system was placed for studies. MRI contrast agent (0.05 ml of 105.8 mg/ml preparation; MultiHance Bracco Diagnostic Inc.) was administered intravenously to confirm tumor location by contrast-enhanced MR imaging. A four-spot grid of sonications was overlaid on the MR-visible tumor and sonications were carried out at a 0.5% duty cycle for 2 minutes. All sonications were performed at a non-derated peak negative pressure of 0.4 MPa. MR imaging was repeated to confirm BBB/BTB disruption. Following FUS exposure, mice were moved to a heating pad and allowed to recover. For repeat BBB/BTB opening procedures, the entirety of this procedure was repeated once every three days for a total of three separate sessions.

### Reagents

All experiments involving the use of radioactive materials were conducted under a radiation use authorization approved by the University of Virginia Radiation Safety Committee. For all studies, radioactivity was determined using a Capintech CRC-15 Dose Calibrator calibrated to a factor of 517 for 89-Zr (Florham Park, NJ). All chemicals were purchased from Millipore-Sigma (St. Louis, MO) unless otherwise specified, and all solutions were prepared using ultrapure water produced by a Millipore water purification system (Millipore, Billerica, MA).

### 89-Zr Production and Purification

[^89^Zr] Zr-oxalic acid is commercially available from 3D Imaging Drug Design and Development, LLC (Little Rock, AR) and was used without subsequent purification for all studies.

### Conjugation and Purification of Antibody

Anti-mouse CD47 (mCD47) (clone MIAP410) antibody was obtained from BioXCell. An aliquot of antibody stock solution was diluted with 0.9% saline to render 2-10 mg/mL concentration for optimal conjugation. The pH was adjusted to 8.9-9.1 with 0.1 M Na_2_CO_3_ (maximum volume of 160 μL). Df-Bz-NCS chelator (3.3 mg) was dissolved in 1 mL of DMSO. Of this solution, 20 μL was added to the antibody solution in 5 μL increments with mixing between each addition. The resulting reaction mixture was incubated at 37 °C for 30 mins while shaking. During this time, a PD-10 column was conditioned with 20 mL of 5 mg/mL Gentisic Acid (in 0.25 M sodium acetate buffer). When the conjugation was complete, the reaction mixture was pipetted onto the PD-10 column and the flow thru was discarded. The PD-10 column was washed with 1.5 mL of 5 mg/mL Gentisic Acid (in 0.25 M sodium acetate buffer) and flow thru was once again discarded. 2 mL of 5 mg/mL Gentisic Acid (in 0.25 M sodium acetate buffer) was then added to the PD-10 column and flow thru was collected in 1 mL fraction. These fractions were stored at -20 °C for no longer than two weeks until labeling took place.

### Radiolabeling of Antibodies

Production of [89Zr]-mCD47 was performed as reported, with minor modifications [49]. A maximum of 200 μL [^89^Zr] Zr-oxalic acid was added to a glass reaction vial equipped with a clean stir bar. While gently stirring, 1 M oxalic acid was added to the reaction vial. 90 μL of 2 M Na_2_CO_3_ was added to the reaction vial and the reaction mixture was stirred at room temperature for 3 mins. Next, 0.3 mL of 0.5 M HEPES, 0.71 mL of the conjugated antibody and 0.7 mL 0.5 M HEPES were added to the reaction vial. The pH was checked for an optimal range of 6.8-7.2. The reaction mixture was incubated at room temperature for 1 hr while stirring. RadioTLC analysis indicated incubations were generally complete by 45 minutes (Rf [89Zr]-mCD47 0.9; mobile phase: 20 mM citric acid, pH 4.9-5.1) During the reaction time, a PD-10 column was conditioned with 20 mL of saline and eluate was discarded. After the reaction was complete, the reaction mixture was added to the PD-10 column and the eluate was discarded. 1.5 mL of saline was added to the PD-10 column and the eluate was discarded. Next, 6 mL of saline was added to the column and six 1 mL fractions were collected. The majority of the labelled antibody was found in the first two fractions. These fractions were used for in vivo injections.

### In Vitro Cell Binding

Approximately 2×10^6^ cells were transferred into microcentrifuge tubes containing sterile PBS and each tube received 1-2 μCi of [^89^Zr]-mCD47. After incubating at 37 °C for 90 minutes, cells (n=5 replicates) were centrifuged at 13,200G for 1 minute and media was removed by vacuum aspiration. Cells were washed 2x with ice-cold PBS and the remaining pellets were analysed using an automated gamma counter (Hidex). Data were reported as becquerels/log 10^6^ cells (Bq/log 10^6^ cells).

### In Vivo Biodistribution

Naïve or tumor-bearing mice received a single intravenous [^89^Zr]-mCD47 injection with or without the accompaniment of FUS BBB/BTB-D. Approximately 3 days later, mice were euthanized and organs were harvested for biodistribution analysis. Brain, liver, spleen, kidneys and whole blood were excised. Brain hemispheres were incised at the midline and collected separately for analysis. Each tissue sample was transferred to a glass scintillation vial and analyzed using an automated gamma counter (Hidex). Following correction for background and decay, the percent injected dose per gram (%ID/g) was calculated by comparison with an aqueous [^89^Zr]oxalate standard.

### Small-Animal PET/CT Imaging

Serial static PET/CT imaging was performed at select time points between 0- and 7-days following intervention with FUS and/or [^89^Zr]-mCD47. Mice previously injected with [^89^Zr]-mCD47 were anesthetized with 1-2% isoflurane and imaged using the Albira Si small-animal trimodal PET/CT/SPECT scanner (Bruker). Whole body static PET images were collected for 20-30 minutes followed by a 10-minute CT scan. PET images were reconstructed using the 3D-OSEM algorithm, with a total of four iterations and 15 subsets, using an isotropic voxel resolution of 0.5×0.5×0.5 mm^3^. CT images were reconstructed using a 3D filtered back projection (FBP) algorithm. Image co-registration and analysis was performed using PMOD 4.1 (PMOD Technologies, Ltd.). First, PET images were co-registered to a CT template, following which VOIs were selected from PET images to encompass either whole brain or each brain hemisphere separately. CT anatomic guidelines were utilized to guide VOI selection. Care was taken to avoid non-cerebral structures and to avoid spillover radioactivity from extracranial structures with high uptake. Averaged standard uptake values (SUVs) were derived for each VOI and data are reported as percent injected dose per mL (%ID/ml).

### Therapeutic mCD47 Delivery

For CD47 blockade therapy, anti-mouse CD47 mAb (MIAP410; BioXCell) diluted in sterilized 0.9% saline was administered intravenously at a dose of either 8 or 32 mg/kg. Control mice received a similarly diluted equivalent dose of mouse IgG1 isotype control antibody (MOPC-21; BioXCell) intravenously. For repeat BBB/BTB-D studies, antibody administration was repeated every three days for a total of three doses, beginning on either day 8 or day 14 post-inoculation. Mice undergoing FUS BBB/BTB-D as described above received anti-CD47 within 15 minutes following conclusion of the procedure.

### Tumor Image Analysis

GL261 tumor outgrowth was monitored via contrast-enhanced MR imaging on Days 14, 17 and 20. Images were analyzed using Horos Viewer (Horos Project). Each MR-visible tumor was traced on all axial MRI slices using the ROI tool, following which a 3D volume was rendered for each tumor. For analysis of tumor permeability, images were analyzed using ImageJ. MR-visible tumor within a single axial MRI slice was traced using a customized ROI for each mouse. Mean greyscale intensity (MGI) was extracted for tumors on Days 17 and 20 and represented as a fold change over background MGI of non-bearing brains.

### Survival Monitoring

After treatment, tumor-bearing mice were monitored for overall survival. Mice were euthanized according to protocol when they displayed physical signs of morbidity (e.g., weight loss, ambulatory deficits, hunched posture) or behavioral deficits indicative of tumor-associated decline.

### Statistical Analysis

All statistical analyses were performed in Graphpad Prism 8 (Graphpad Software, Inc). Data are shown as mean ± SEM unless otherwise noted. A detailed description of statistical methods for each experiment is provided in the corresponding figure legend. Individual data points are provided in all data figures, thus “n” values per group are always represented.

## Data Availability

The authors declare that [the/all other] data supporting the findings of this study are available within this paper and is supplementary information files.

## Acknowledgements

The authors thank Dr. Sasha Klibanov (UVA) for supplying microbubbles for all FUS studies, as well as Mr. Jeremy Gatesman and Mr. Jose Reyes for their invaluable assistance with study planning and technical execution of radiolabeled antibody delivery experiments. This study was supported by National Institutes of Health Grants R01CA197111, R01EB020147 and R21NS118278 to R.J.P. N.D.S. was supported by a National Cancer Institute F99/K00 Predoctoral to Postdoctoral Fellow Transition Award (F99CA234954), NSF Graduate Research Fellowship, and UVA School of Medicine Wagner Fellowship.

## Author Contributions

N.D.S., S.P., K.S.M. and K.D.N. conceived and performed key experiments and analyzed the data. V.B. performed MR image analysis. G.W.M. developed MR sequences and performed MR imaging. S.S.B. contributed to experiment design and supervised studies. R.J.P. provided funding support and supervised all studies. N.D.S. and R.J.P. wrote the manuscript. All authors discussed results and contributed to completion of the paper.

## Competing Interests

The authors declare no competing interests.

**Correspondence** and requests for materials should be addressed to N.D.S.

## References

1. Stupp R, Mason WP, Van Den Bent MJ, Weller M, Fisher B, Taphoorn MJB, Belanger K, Brandes AA, Marosi C, Bogdahn U. Radiotherapy plus concomitant and adjuvant temozolomide for glioblastoma. N Engl J Med. 2005; 352:987–996.

2. Razavi S-M, Lee KE, Jin BE, Aujla PS, Gholamin S, Li G. Immune evasion strategies of glioblastoma. Front Surg. 2016; 3:11.

3. Gholamin S, Mitra SS, Feroze AH, Liu J, Kahn SA, Zhang M, Esparza R, Richard C, Ramaswamy V, Remke M, Volkmer AK, Willingham S, Ponnuswami A, McCarty A, Lovelace P, Storm TA, Schubert S, Hutter G, Narayanan C, Chu P, Raabe EH, Harsh G, Taylor MD, Monje M, Cho YJ, Majeti R, Volkmer JP, Fisher PG, Grant G, Steinberg GK, Vogel H, Edwards M, Weissman IL, Cheshier SH. Disrupting the CD47-SIRPα anti-phagocytic axis by a humanized anti-CD47 antibody is an efficacious treatment for malignant pediatric brain tumors. Sci Transl Med. 2017; 9:1–14.

4. Hutter G, Theruvath J, Graef CM, Zhang M, Schoen MK, Manz EM, Bennett ML, Olson A, Azad TD, Sinha R. Microglia are effector cells of CD47-SIRPα antiphagocytic axis disruption against glioblastoma. Proc Natl Acad Sci. 2019; 116:997–1006.

5. Zhang M, Hutter G, Kahn SA, Azad TD, Gholamin S, Xu CY, Liu J, Achrol AS, Richard C, Sommerkamp P, Schoen MK, McCracken MN, Majeti R, Weissman I, Mitra SS, Cheshier SH. Anti-CD47 Treatment Stimulates Phagocytosis of Glioblastoma by M1 and M2 Polarized Macrophages and Promotes M1 Polarized Macrophages In Vivo. PLoS One. 459 2016; 11:e0153550.

6. Zhao H-J, Pan F, Shi Y-C, Luo X, Ren R-R, Peng L-H, Yang Y-S. Prognostic significance of CD47 in human malignancies: a systematic review and meta-analysis. Trans Cancer Res. 2018; 7:609–621.

7. Barclay AN, van den Berg TK. The Interaction Between Signal Regulatory Protein Alpha (SIRP α) and CD47: Structure, Function, and Therapeutic Target. Annu Rev Immunol. 2014; 32:25–50.

8. Willingham SB, Volkmer J-P, Gentles AJ, Sahoo D, Dalerba P, Mitra SS, Wang J, Contreras-Trujillo H, Martin R, Cohen JD, Lovelace P, Scheeren FA, Chao MP, Weiskopf K, Tang C, Volkmer AK, Naik TJ, Storm TA, Mosley AR, Edris B, Schmid SM, Sun CK, Chua M-S, Murillo O, Rajendran P, Cha AC, Chin RK, Kim D, Adorno M, Raveh T, Tseng D, Jaiswal S, Enger PØ, Steinberg GK, Li G, So SK, Majeti R, Harsh GR, van de Rijn M, Teng NNH, Sunwoo JB, Alizadeh AA, Clarke MF, Weissman IL. The CD47-signal regulatory protein alpha (SIRPa) interaction is a therapeutic target for human solid tumors. Proc Natl Acad Sci U S A. 2012; 109:6662–7.

9. Li F, Lv B, Liu Y, Hua T, Han J, Sun C, Xu L, Zhang Z, Feng Z, Cai Y, Zou Y, Ke Y, Jiang X. Blocking the CD47-SIRPα axis by delivery of anti-CD47 antibody induces antitumor effects in glioma and glioma stem cells. Oncoimmunology. 2017; 7:e1391973–e1391973.

10. Chao MP, Jaiswal S, Weissman-Tsukamoto R, Alizadeh AA, Gentles AJ, Volkmer J, Weiskopf K, Willingham SB, Raveh T, Park CY, Majeti R, Weissman IL. Calreticulin Is the Dominant Pro-Phagocytic Signal on Multiple Human Cancers and Is Counterbalanced by CD47. Sci Transl Med. 2010; 2:63ra94–63ra94.

11. von Roemeling CA, Wang Y, Qie Y, Yuan H, Zhao H, Liu X, Yang Z, Yang M, Deng W, Bruno KA, Chan CK, Lee AS, Rosenfeld SS, Yun K, Johnson AJ, Mitchell DA, Jiang W, Kim BYS. Therapeutic modulation of phagocytosis in glioblastoma can activate both innate and adaptive antitumour immunity. Nat Commun. 2020; 11:1508.

12. Chacko A-M, Li C, Pryma DA, Brem S, Coukos G, Muzykantov V. Targeted delivery of antibody-based therapeutic and imaging agents to CNS tumors: crossing the blood-brain barrier divide. Expert Opin Drug Deliv. 2013; 10:907–926.

13. Razpotnik R, Novak N, Čurin Šerbec V, Rajcevic U. Targeting Malignant Brain Tumors with Antibodies. Front Immunol. 2017; 8:1181.

14. Patel M, McCully C, Godwin K, Balis FM. Plasma and cerebrospinal fluid pharmacokinetics of intravenous temozolomide in non-human primates. J Neurooncol. 2003; 61:203–207.

15. Rosso L, Brock CS, Gallo JM, Saleem A, Price PM, Turkheimer FE, Aboagye EO. A new model for prediction of drug distribution in tumor and normal tissues: pharmacokinetics of temozolomide in glioma patients. Cancer Res. 2009; 69:120–127.

16. van Tellingen O, Yetkin-Arik B, de Gooijer MC, Wesseling P, Wurdinger T, de Vries HE. Overcoming the blood–brain tumor barrier for effective glioblastoma treatment. Drug Resist Updat. 2015; 19:1–12.

17. Timbie KF, Mead BP, Price RJ. Drug and gene delivery across the blood-brain barrier with focused ultrasound. J Control Release. 2015; 219:61–75.

18. Kinoshita M, McDannold N, Jolesz FA, Hynynen K. Noninvasive localized delivery of Herceptin to the mouse brain by MRI-guided focused ultrasound-induced blood-brain barrier disruption. ProcNatlAcadSciUSA. no date; 103:11719–11723.

19. Arvanitis CD, Askoxylakis V, Guo Y, Datta M, Kloepper J, Ferraro GB, Bernabeu MO, Fukumura D, McDannold N, Jain RK. Mechanisms of enhanced drug delivery in brain metastases with focused ultrasound-induced blood-tumor barrier disruption. Proc Natl Acad Sci U S A. 2018; 115:E8717–E8726.

20. Liu H-L, Hsu P-H, Lin C-Y, Huang C-W, Chai W-Y, Chu P-C, Huang C-Y, Chen P-Y, Yang L-Y, Kuo JS, Wei K-C. Focused Ultrasound Enhances Central Nervous System Delivery of Bevacizumab for Malignant Glioma Treatment. Radiology. 2016; 281:99–108.

21. Park EJ, Zhang YZ, Vykhodtseva N, McDannold N. Ultrasound-mediated blood-brain/blood-tumor barrier disruption improves outcomes with trastuzumab in a breast cancer brain metastasis model. J Control Release. 2012; 163:277–284.

22. Lipsman N, Meng Y, Bethune AJ, Huang Y, Lam B, Masellis M, Herrmann N, Heyn C, Aubert I, Boutet A, Smith GS, Hynynen K, Black SE. Blood–brain barrier opening in Alzheimer’s disease using MR-guided focused ultrasound. Nat Commun. 2018; 9:1–8.

23. Abrahao A, Meng Y, Llinas M, Huang Y, Hamani C, Mainprize T, Aubert I, Heyn C, Black SE, Hynynen K, Lipsman N, Zinman L. First-in-human trial of blood-brain barrier opening in amyotrophic lateral sclerosis using MR-guided focused ultrasound. Nat Commun. 2019; 10:4373.

24. Mainprize T, Lipsman N, Huang Y, Meng Y, Bethune A, Ironside S, Heyn C, Alkins R, Trudeau M, Sahgal A, Perry J, Hynynen K. Blood-Brain Barrier Opening in Primary Brain Tumors with Non-invasive MR-Guided Focused Ultrasound: A Clinical Safety and Feasibility Study. Sci Rep. 2019; 9:321.

25. Timbie KFK, Afzal U, Date A, Zhang C, Song J, Wilson Miller G, Suk JJS, Hanes J, Price RJR, Miller G, Suk JJS, Hanes J, Price RJR. MR Image-Guided Delivery of Cisplatin-Loaded Brain-Penetrating Nanoparticles to Invasive Glioma with Focused Ultrasound. J Control Release. 2017; 263:120–131.

26. Park J, Aryal M, Vykhodtseva N, Zhang Y-Z, McDannold N. Evaluation of permeability, doxorubicin delivery, and drug retention in a rat brain tumor model after ultrasound-induced blood-tumor barrier disruption. J Control Release. 2017; 250:77–85.

27. Park J, Zhang Y, Vykhodtseva N, Jolesz FA, McDannold NJ. The kinetics of blood brain barrier permeability and targeted doxorubicin delivery into brain induced by focused ultrasound. J Control Release. 2012; 162:134–142.

28. Kinoshita M, McDannold N, Jolesz FA, Hynynen K. Noninvasive localized delivery of Herceptin to the mouse brain by MRI-guided focused ultrasound-induced blood-brain barrier disruption. Proc Natl Acad Sci U S A. 2006; 103:11719–11723.

29. Kinoshita M, McDannold N, Jolesz FA, Hynynen K. Targeted delivery of antibodies through the blood–brain barrier by MRI-guided focused ultrasound. Biochem Biophys Res Commun. 2006; 340:1085–1090.

30. Raymond SB, Treat LH, Dewey JD, McDannold NJ, Hynynen K, Bacskai BJ. Ultrasound enhanced delivery of molecular imaging and therapeutic agents in Alzheimer’s disease mouse models. PLoS One. 2008; 3:1–7.

31. Jordão JF, Ayala-Grosso CA, Markham K, Huang Y, Chopra R, McLaurin J, Hynynen K, Aubert I. Antibodies targeted to the brain with image-guided focused ultrasound reduces amyloid-β plaque load in the TgCRND8 mouse model of Alzheimer’s disease. PLoS One. 2010; 5:4–11.

32. Jordão JF, Thévenot E, Markham-Coultes K, Scarcelli T, Weng YQ, Xhima K, O’Reilly M, Huang Y, McLaurin J, Hynynen K, Aubert I. Amyloid-β plaque reduction, endogenous antibody delivery and glial activation by brain-targeted, transcranial focused ultrasound. Exp Neurol. 2013; 248:16–29.

33. Nisbet RM, Van der Jeugd A, Leinenga G, Evans HT, Janowicz PW, Götz J. Combined effects of scanning ultrasound and a tau-specific single chain antibody in a tau transgenic mouse model. Brain. 2017; 1–11.

34. Kobus T, Zervantonakis IK, Zhang Y, McDannold NJ. Growth inhibition in a brain metastasis model by antibody delivery using focused ultrasound-mediated blood-brain barrier disruption. J Control Release. 2016; 238:281–288.

35. Carpentier A, Canney M, Vignot A, Reina V, Beccaria K, Horodyckid C, Karachi C, Leclercq D, Lafon C, Chapelon J-Y, Capelle L, Cornu P, Sanson M, Hoang-Xuan K, Delattre J-Y, Idbaih A. Clinical trial of blood-brain barrier disruption by pulsed ultrasound. Sci Transl Med. 2016; 8:343re2.

36. Mead BP, Mastorakos P, Suk JS, Klibanov AL, Hanes J, Price RJ. Targeted gene transfer to the brain via the delivery of brain-penetrating DNA nanoparticles with focused ultrasound. J Control Release. 2016; 223:109–117.

37. Curley CT, Mead BP, Negron K, Garrison WJ, Miller GW, Kingsmore KM, Thim EA, Song J, Munson JM, Klibanov AL, Suk JS, Hanes J, Price RJ. Augmentation of Brain Tumor Interstitial Flow via Focused Ultrasound Promotes Brain-Penetrating Nanoparticle Dispersion and Transfection. Sci Adv. 2020; 6:aay1344.

38. Aryal M, Vykhodtseva N, Zhang Y-Z, Park J, McDannold N. Multiple treatments with liposomal doxorubicin and ultrasound-induced disruption of blood-tumor and blood-brain barriers improve outcomes in a rat glioma model. J Control Release. 2013; 169:103–11.

39. Kobus T, Zervantonakis IK, Zhang Y, McDannold NJ. Growth inhibition in a brain metastasis model by antibody delivery using focused ultrasound-mediated blood-brain barrier disruption. J Control Release. 2016; 238:281–288.

40. Chao MP, Weissman IL, Majeti R. The CD47–SIRPα pathway in cancer immune evasion and potential therapeutic implications. Curr Opin Immunol. 2012; 24:225–232.

41. Cousin S, Seneschal J, Italiano A. Toxicity profiles of immunotherapy. Pharmacol Ther. 2018; 181:91–100.

42. O’Reilly MA, Hough O, Hynynen K. Blood-Brain Barrier Closure Time After Controlled Ultrasound-Induced Opening Is Independent of Opening Volume. J Ultrasound Med. 2017;.

43. Gao L, Chen K, Gao Q, Wang X, Sun J, Yang Y-G. CD47 deficiency in tumor stroma promotes tumor progression by enhancing angiogenesis. Oncotarget. 2017; 8:22406–22413.

44. Kaur S, Martin-Manso G, Pendrak ML, Garfield SH, Isenberg JS, Roberts DD. Thrombospondin-1 inhibits VEGF receptor-2 signaling by disrupting its association with CD47. J Biol Chem. 2010; 285:38923–38932.

45. Aryal M, Arvanitis CD, Alexander PM, McDannold N. Ultrasound-mediated blood–brain barrier disruption for targeted drug delivery in the central nervous system. Adv Drug Deliv Rev. 2014; 72:94–109.

46. Arif WM, Elsinga PH, Gasca-Salas C, Versluis M, Martínez-Fernández R, Dierckx RAJO, Borra RJH, Luurtsema G. Focused ultrasound for opening blood-brain barrier and drug delivery monitored with positron emission tomography. J Control Release. 2020;.

47. Rezai AR, Ranjan M, D’Haese P-F, Haut MW, Carpenter J, Najib U, Mehta RI, Chazen JL, Zibly Z, Yates JR, Hodder SL, Kaplitt M. Noninvasive hippocampal blood-brain barrier opening in Alzheimer’s disease with focused ultrasound. Proc Natl Acad Sci U S A. 2020; 117:9180–9182.

48. Zhang W, Huang Q, Xiao W, Zhao Y, Pi J, Xu H, Zhao H, Xu J, Evans CE, Jin H. Advances in Anti-Tumor Treatments Targeting the CD47/SIRPα Axis. Front Immunol. 2020; 11:18.

49. Zheleznyak A, Ikotun OF, Dimitry J, Frazier WA, Lapi SE. Imaging of CD47 expression in xenograft and allograft tumor models. Mol Imaging. 2013; 12:1–10.

