## Supplemental Figures for "ImmunoPET-informed sequence for focused ultrasound-targeted mCD47 blockade controls glioma"

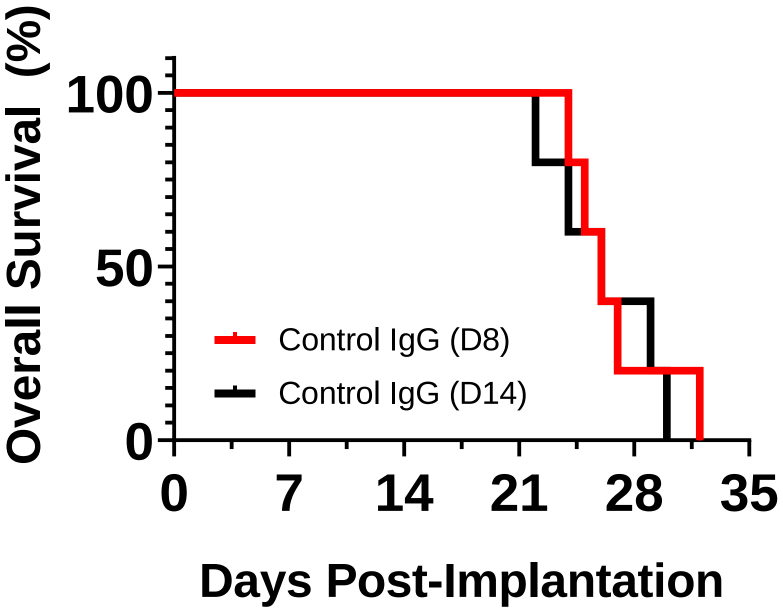


**Supplemental Figure 1. Overall survival of control GL261-bearing mice.** Kaplan-Meier curve depicting overall survival of control GL261-bearing mice receiving systemic control IgG antibody. Treatments were initiated either 8 (D8) or 14 (D14) days post-implantation. *n*=5 mice per group. Significance assessed by log-rank (Mantel-Cox) test. No significant difference detected.

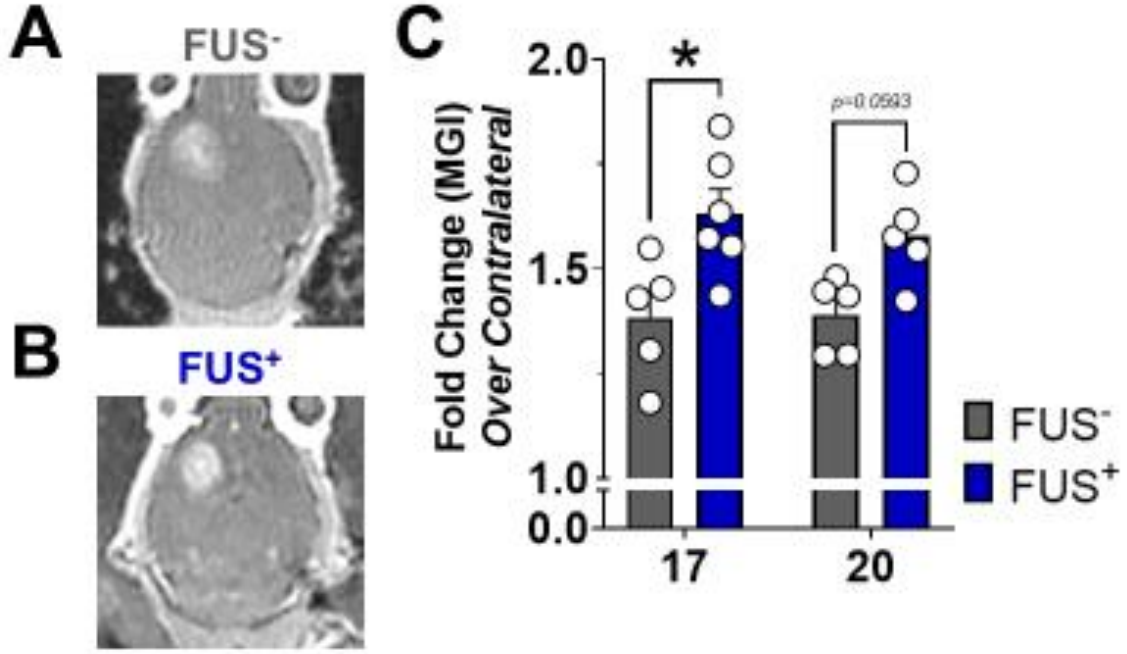


**Supplemental Figure 2. BBB/BTB permeability in baseline pre-treatment planning MRIs from mCD47 survival study.** (A, B) Representative baseline contrast-enhanced MR images of GL261 tumor-bearing brains at Day 17. (C) Mean greyscale intensity (MGI) of contrast enhancement for GL261 tumors in $FUS^-$ and $FUS^+$ groups, calculated as fold change over contralateral brain. *p=0.0093. Significance assessed by RM mixed effects model implementing restricted maximum likelihood method, followed by Sidak's multiple comparison correction.
